# Molecular determinants of extracellular TIMP-3 accumulation in Sorsby fundus dystrophy

**DOI:** 10.1101/2025.05.19.654781

**Authors:** Jacob H. J. Betts, Katherine Hampton, Dudley K. Strickland, Simon J. Clark, Anthony J. Day, Linda Troeberg

## Abstract

Tissue inhibitor of metalloproteinases-3 (TIMP-3) is a critical regulator of extracellular matrix turnover. Mutations in TIMP-3 cause Sorsby fundus dystrophy (SFD), an inherited macular dystrophy that is similar to age-related macular degeneration (AMD) but which generally presents earlier. SFD is characterised by the accumulation of mutant TIMP-3 protein in Bruch’s membrane, a multilaminar extracellular matrix underlying the retinal pigment epithelium (RPE). Here, we show that RPE cells regulate wild-type TIMP-3 levels post-translationally, with ARPE-19 and hTERT RPE-1 cell lines endocytosing the protein via the low-density lipoprotein receptor-related protein (LRP) family of scavenger receptors. LRP-mediated endocytosis of the SFD TIMP-3 variants S204C and Y191C was significantly delayed, establishing a molecular mechanism for their extracellular accumulation in SFD. In contrast, endocytosis of the SFD variant H181R TIMP-3 was unaltered, suggesting it accumulates through a distinct molecular mechanism, potentially via increased retention on extracellular matrix heparan sulfate proteoglycans. These findings reveal heterogeneity in the molecular mechanism of SFD pathogenesis, which has direct implications for therapeutic development. Genotype-specific interventions may be required, such as strategies that enhance receptor-mediated clearance or disrupt extracellular TIMP-3 retention. Our study also has broader implications for AMD, where TIMP-3 and other LRP and heparan sulfate ligands accumulate in drusen within Bruch’s membrane. Age- and inflammation-dependent alterations in LRP expression and heparan sulfate structure may contribute to drusen formation and AMD progression. Understanding TIMP-3 trafficking in both physiological and pathological contexts could inform targeted treatments for SFD, AMD, and other degenerative disorders involving extracellular matrix dysregulation.

## Introduction

Sorsby fundus dystrophy (SFD) is an early-onset, inherited form of retinal degeneration caused by autosomal dominant mutations in the coding region of the *TIMP3* gene, which encodes tissue inhibitor of metalloproteinases 3 (TIMP-3)(1). First identified by Sorsby and Mason in 1949 (2), SFD shares many clinical features with age-related macular degeneration (AMD), which is a leading cause of sight loss worldwide (3). While AMD has complex genetic and environmental causes, the known genetic aetiology of SFD makes it a tractable model in which to define molecular mechanisms modulating retinal health and degeneration.

The American College of Genetics and Genomics (ACMG) and the Association for Molecular Pathology (AMP) guidelines classify 11 known *TIMP3* mutations as pathogenic for SFD and another 11 *TIMP3* mutations as likely to be pathogenic (4). Most (4) of these mutations (17/22) result in an unpaired cysteine residue, which is thought to disrupt disulfide bond formation. SFD variants of TIMP-3 accumulate extracellularly in deposits or “drusen” within the Bruch’s membrane, a five-layer extracellular matrix (ECM) underlying the retinal pigment epithelium (RPE). Similar drusen form in AMD, where the Bruch’s membrane contains elevated levels of a variety of lipids and proteins (5), including TIMP-3 (6, 7). Drusen formation is associated with thickening of the Bruch’s membrane (8), along with macular atrophy and/or neovascularization that progressively impairs vision.

Various molecular mechanisms have been proposed to explain how extracellular accumulation of SFD TIMP-3 mutants causes retinal pathology (9, 10), although these mechanisms have been studied for only a handful of SFD TIMP-3 mutations and may not apply equally to all. Firstly, while affinity of SFD TIMP-3 mutants for prototypic matrix metalloproteinases (MMPs) is not substantially affected, overall inhibition of metalloproteinases is likely to be increased in the Bruch’s membrane due to increased extracellular accumulation of mutant TIMP-3. This is proposed to reduce turnover of extracellular matrix (ECM) components and hence to cause thickening of the Bruch’s membrane (11). Secondly, TIMP-3 has been shown to induce apoptosis of cancer cells, through inhibition of adamalysins that shed death receptors (12, 13). TIMP-3 has also been reported to induce apoptosis of RPE cells, with SFD TIMP-3 mutants reducing cell viability significantly more than WT TIMP-3 (14). TIMP-3 inhibits angiogenesis through its ability to inhibit VEGFR signaling (15), and SFD mutants of TIMP-3 are reported to inhibit VEGFR signaling less effectively (16, 17) and so to contribute to the neovascularization seen in SFD. More recently, altered metabolism has been observed in RPE cells expressing SFD TIMP-3 mutants (18, 19). It is not clear how mutated TIMP-3 causes these changes, but such metabolic dysfunction could contribute to retinal damage.

Whatever the mechanism(s) are by which accumulated SFD TIMP-3 mutants damage the retina, a key unanswered question is what causes their increased extracellular abundance. TIMP-3 mRNA levels are not increased in SFD (20), suggesting that the mutations alter post-translational regulation of the protein. This hypothesis is supported by investigations by Langton et al., who found that SFD mutants of TIMP-3 (E162X, S179C, S204C) were cleared from the extracellular environment more slowly than WT TIMP-3, leading them to propose that SFD mutants are “resistant to turnover” (21). We subsequently found that fibroblasts, chondrosarcoma cells and macrophages can endocytose WT TIMP-3 via the scavenger receptor low-density lipoprotein receptor-related protein 1 (LRP1)(22–24). This endocytosis is inhibited by heparan sulfate proteoglycans (HSPGs) in the extracellular matrix (ECM) (22, 25), suggesting that extracellular TIMP-3 abundance is determined by the equilibrium between its binding to LRP1 and to heparan sulfate (HS).

Here, we set out to establish whether RPE cells endocytose TIMP-3 and if so, whether this occurs via LRP1. Secondly, we compared endocytosis of WT TIMP-3 to that of the SFD TIMP-3 mutants S204C, Y191C and H181R. These experiments shed light on the molecular mechanisms of SFD pathogenesis, and identify the LRP family of receptors as important mediators of retinal protein turnover and health.

## Results

### Generation of SFD TIMP-3 mutants

Site-directed mutagenesis was used to generate the SFD TIMP-3 mutants H181R, S204C, and Y191C bearing a C-terminal FLAG-tag, and wild-type (WT) and mutant TIMP-3 were isolated from conditioned media of transiently transfected HEK-293-EBNA cells by anti-FLAG M2 affinity chromatography (**Fig. 1A**).

**Fig. 1:**
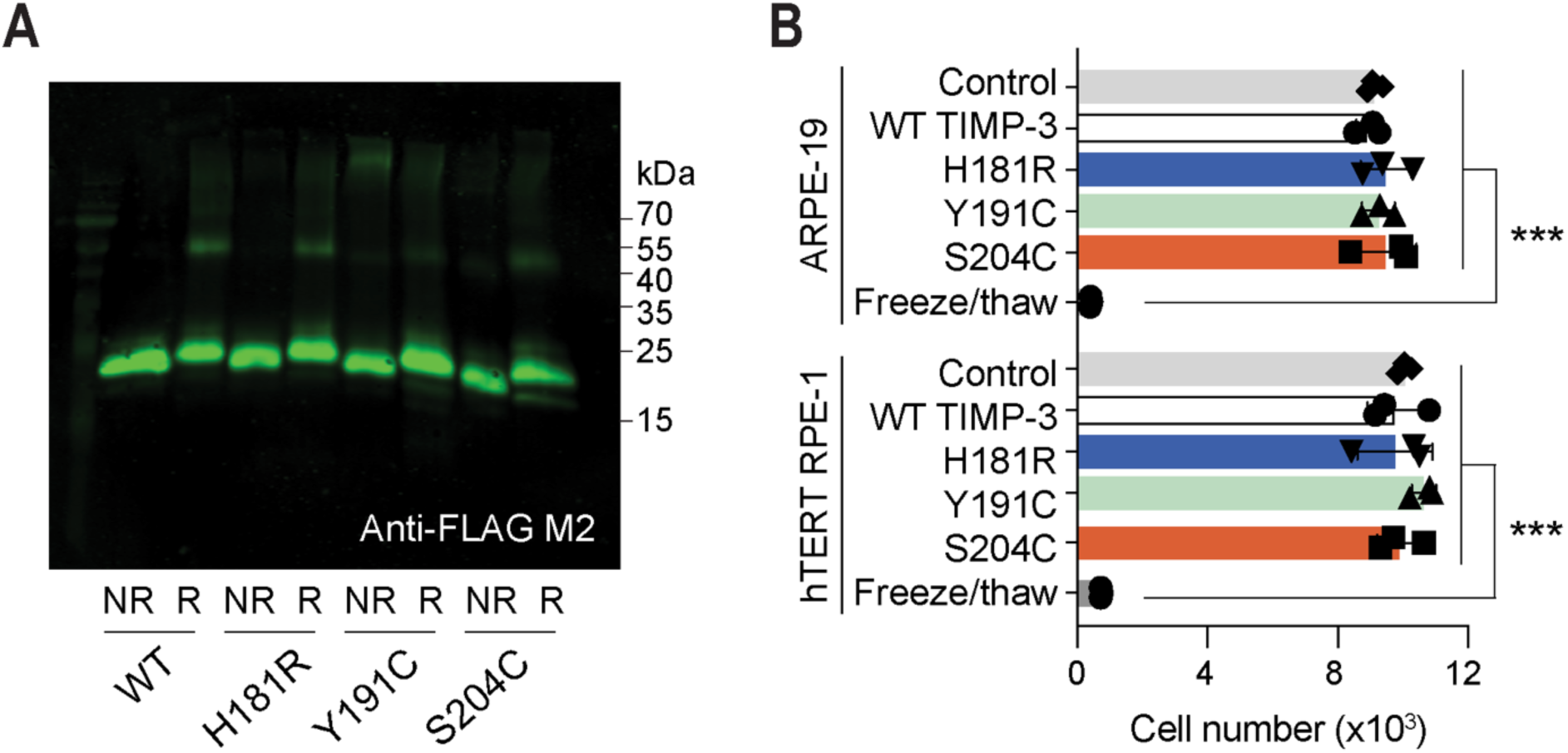
Isolation of WT and SFD mutants of TIMP-3. **A)** Isolated FLAG-tagged WT, H181R, Y191C and S204C TIMP-3 (25 ng each) were either mixed with SDS-PAGE sample buffer (non-reducing, NR), or reduced and alkylated with β-mercaptoethanol and iodoacetamide before addition of SDS-PAGE sample buffer (reducing, R). Samples were electrophoresed on a 15% Tris-glycine SDS-PAGE gel and analysed by immunoblotting with an anti-FLAG M2 primary antibody and a IRDye 800CW goat anti-mouse secondary antibody. **B)** ARPE-19 and hTERT RPE-1 cells (1×10^4^) were incubated (24 h) with WT or SFD TIMP-3 [2.5 nM in DMEM with 0.2% (v/v) FBS] before quantification of cell number with MTS reagent (mean ± SD, n = 3). As a control for cell death, cells were frozen before quantification. Data within each cell type were assessed for normality by Shapiro-Wilk test, analysed for significance by one-way ANOVA, and corrected for multiple comparisons with Tukey’s test (*** p ≤ 0.001).

Addition of the recombinant proteins (2.5 nM) to ARPE-19 and hTERT RPE-19 cells for 24 h had no effect on cell viability as assessed by MTS assay (**Fig. 1B**). Even at a higher concentration (100 nM), there was no evidence of apoptosis in ARPE-19 cells treated with the recombinant proteins for 16 h (**Fig. 2**).

**Fig. 2:**
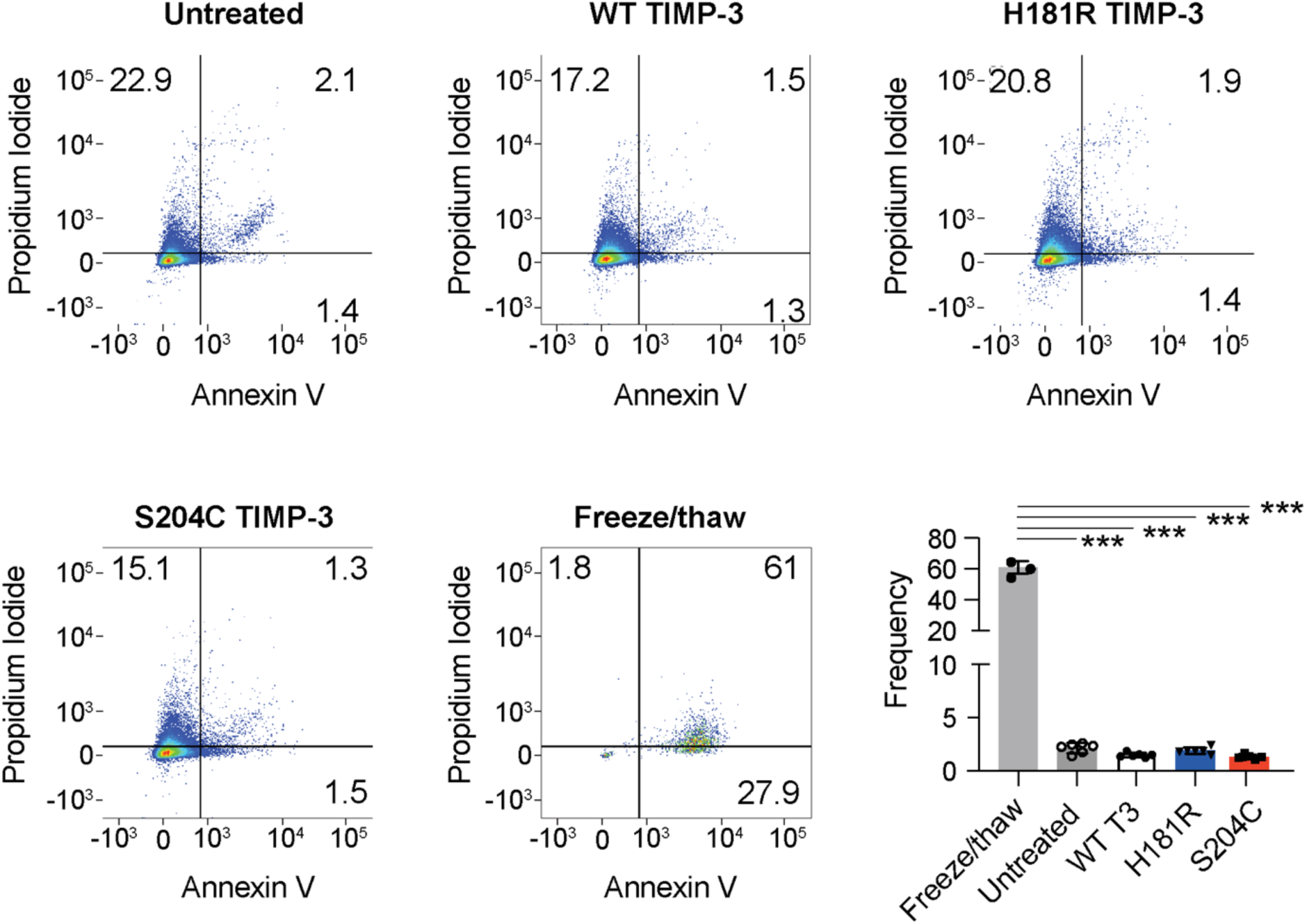
No apoptosis was detected in ARPE-19 cells treated with WT or SFD TIMP-3. ARPE-19 cells (2.5×10^5^) were treated with FLAG-tagged WT, H181R or S204C TIMP-3 (100 nM in serum-free DMEM) or no TIMP-3 (untreated) for 16 h. A positive control for cell death was generated by repeated freeze-thawing. Cells were stained with annexin V and propidium iodide and analysed using flow cytometry. Representative plots indicating the percentage of cells in each quadrant are shown, along with the frequency of apoptotic cells (mean ± SD, n=3 for freeze/thaw, n=6 for other conditions). Data were assessed for normality by Shapiro-Wilk test, analysed for significance with a one-way ANOVA, and corrected for multiple comparisons with Tukey’s test. The only significant difference was between the positive control and all other groups (*** p ≤ 0.001).

### RPE cells cleared extracellular WT TIMP-3 more rapidly than the SFD variants S204C and Y191C

To evaluate cell-mediated clearance of extracellular WT and SFD TIMP-3, we added recombinant FLAG-tagged TIMP-3 WT, H181R, Y191C and S204C to the medium of monolayer cultures of ARPE-19 and hTERT RPE-1 cells, and quantified TIMP-3 abundance in the media over time by anti-FLAG M2 immunoblotting.

This showed that WT TIMP-3 levels in the conditioned media of ARPE-19 cells reduced over 24 h (**Fig. 3A-B**), with a calculated half-life of 2.6 ± 1.1 h. This rate of clearance is similar to that we previously observed in similar experiments with HTB94 chondrosarcoma cells, fibroblasts, and macrophages (23, 24, 26).

**Fig. 3:**
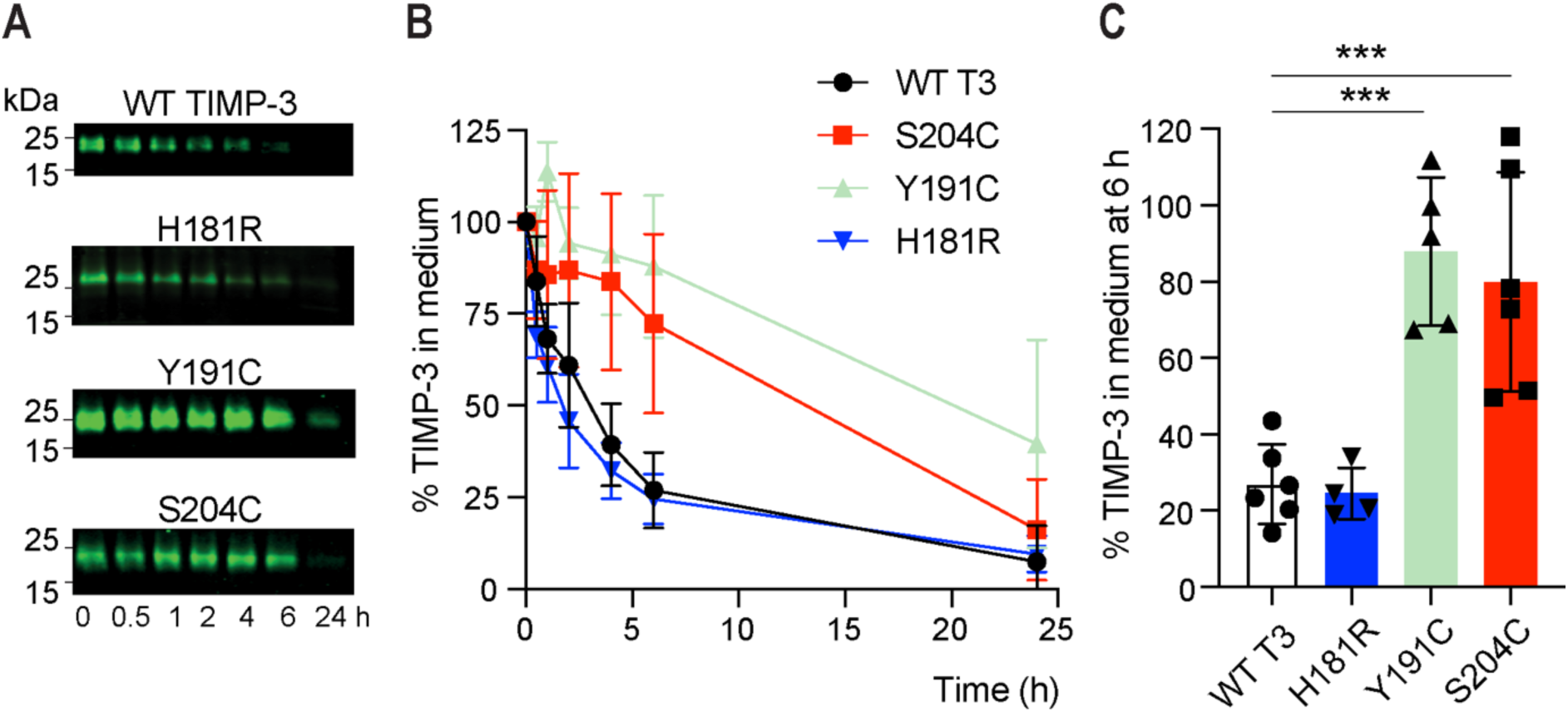
ARPE-19 cells endocytosed WT TIMP-3 and H181R more rapidly than the S204C and Y191C variants. ARPE-19 cells (2.5×10^5^) were incubated (0-24 h) with FLAG-tagged TIMP-3 WT, H181R, Y191C or S204C (2.5 nM) in DMEM with 0.2% (v/v) FBS. **A)** Conditioned media were harvested at the indicated timepoints, concentrated by TCA precipitation, and TIMP-3 abundance quantified by immunoblotting with anti-FLAG M2 antibody and densitometry. Representative immunoblots are shown. **B)** Loss of TIMP-3 variants from conditioned media was calculated relative to total protein staining per lane and to mean pixel volume at t = 0 h (defined as 100%, n = 4-6, mean ± SD). **C)** The percentage of each TIMP-3 variant remaining in conditioned media at 6 h was calculated, with data assessed for normality by Shapiro-Wilk test, analysed for significance relative to WT TIMP-3 with one-way ANOVA, and corrected for multiple comparisons with Tukey’s test (*** p ≤ 0.001).

In contrast, the SFD variants S204C and Y191C were cleared from conditioned media more slowly. A significantly longer half-life (13.9 ± 7.2 h, p ≤ 0.01) was calculated for S204C, while no half-life could be calculated for Y191C as more than 50% of the protein remained in media at 24 h in several experiments (**Fig. 3B**). We therefore compared abundance of the variants in the conditioned media at 6 h, and found that significantly more S204C and Y191C remained in media at this timepoint in comparison with WT TIMP-3 (**Fig. 3C**).

H181R TIMP-3 was cleared from the conditioned media of ARPE-19 cells with the same kinetics as WT TIMP-3 (**Fig. 3A-C**, calculated half-life 1.5 ± 0.8 h), pointing to a marked heterogeneity in the molecular mechanisms that regulate the extracellular accumulation of individual SFD TIMP-3 mutants.

WT and SFD TIMP-3 variants accumulated in vesicle-like structures in ARPE-19 cells, consistent with endocytic uptake (**Fig. 4**, with negative controls in Supplementary **Fig. S1**). A similar pattern of clearance from the extracellular environment was observed with the hTERT RPE-1 cell line (**Supplementary Fig. S2,** WT TIMP-3 half-life 4.9 ± 2.5 h, H181R half-life 5.0 ± 1.7 h).

**Fig. 4:**
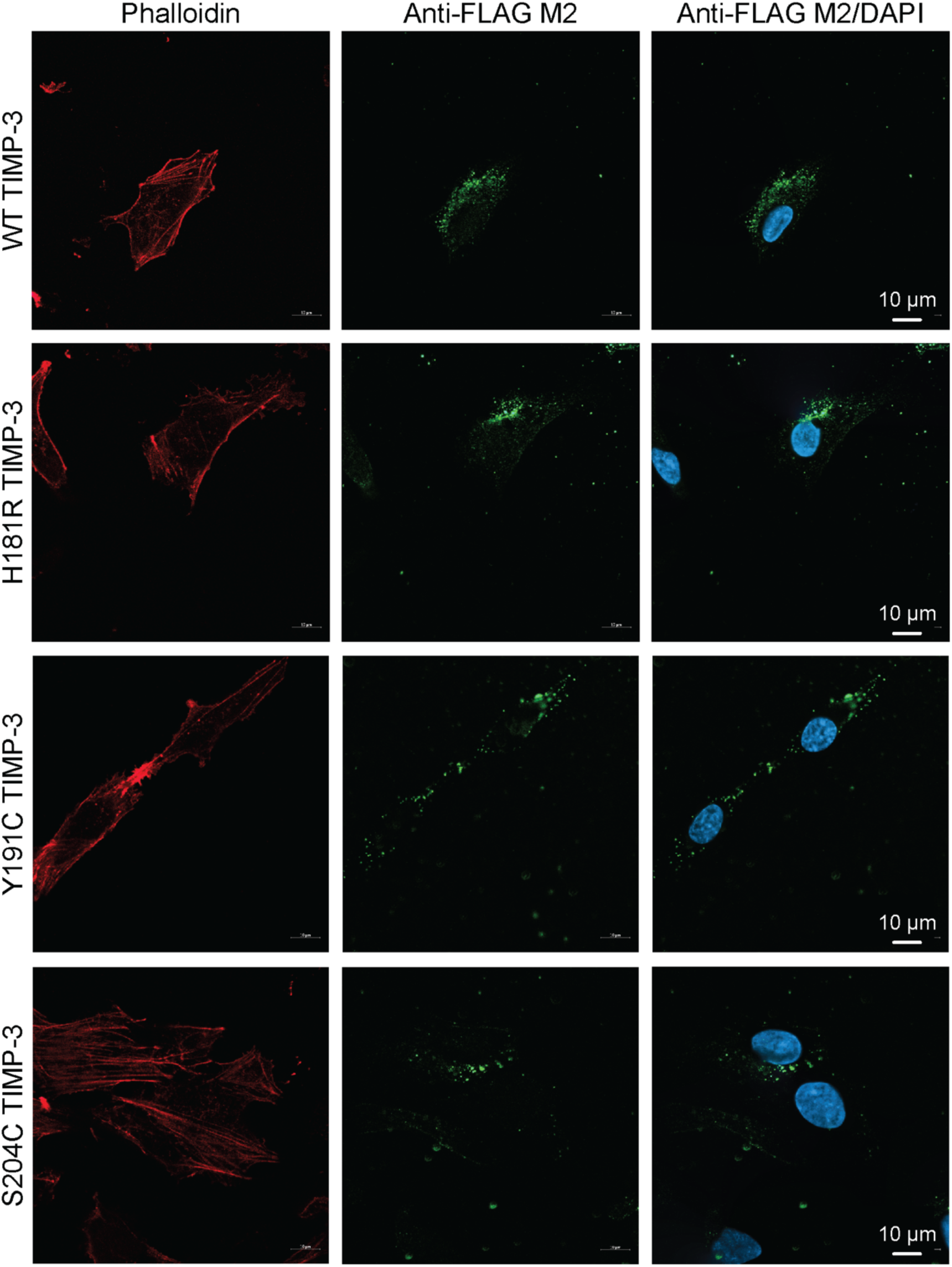
WT and SFD TIMP-3 were visible within ARPE-19 cells. ARPE-19 cells (1×10^5^) were seeded on gelatin-coated coverslips, and treated with FLAG-tagged WT, H181R, Y191C or S204C TIMP-3 (40 nM) in DMEM with 0.2% (v/v) FBS for either 2 h (for WT and H181R TIMP-3) or 18 h (for Y191C and S204C TIMP-3). Cells were permeabilised and stained with anti-FLAG M2 primary antibody and Alexa Fluor-488-conjugated goat anti-mouse IgG. The actin cytoskeleton was stained with Alexa Fluor-647-conjugated phalloidin and nuclei stained with DAPI.

### The LRP family antagonist RAP blocked clearance of extracellular WT and SFD TIMP-3 by RPE cells

Given that LRP1 mediates uptake of WT TIMP-3 in other cells types (23, 24, 26), we investigated whether receptor-associated protein (RAP), an antagonist of ligand binding to the LRP family of receptors, had any effect on clearance of TIMP-3 by RPE cells. Levels of WT and H181R TIMP-3 in conditioned media were significantly reduced after 6 h of incubation with ARPE-19 cells, but no significant clearance was evident when cells were pre-incubated with RAP (**Fig. 5A-B**). Similarly, levels of Y191C and S204C were significantly reduced after 24 h of incubation with ARPE-19 cells, but pre-incubation of the cells with RAP abolished this clearance (**Fig. 5C-D**). RAP also blocked clearance of WT TIMP-3 by hTERT RPE-1 cells (**Supplementary Fig. S3**), indicating that an LRP family member is responsible for endocytosis by both RPE cell lines.

**Fig. 5:**
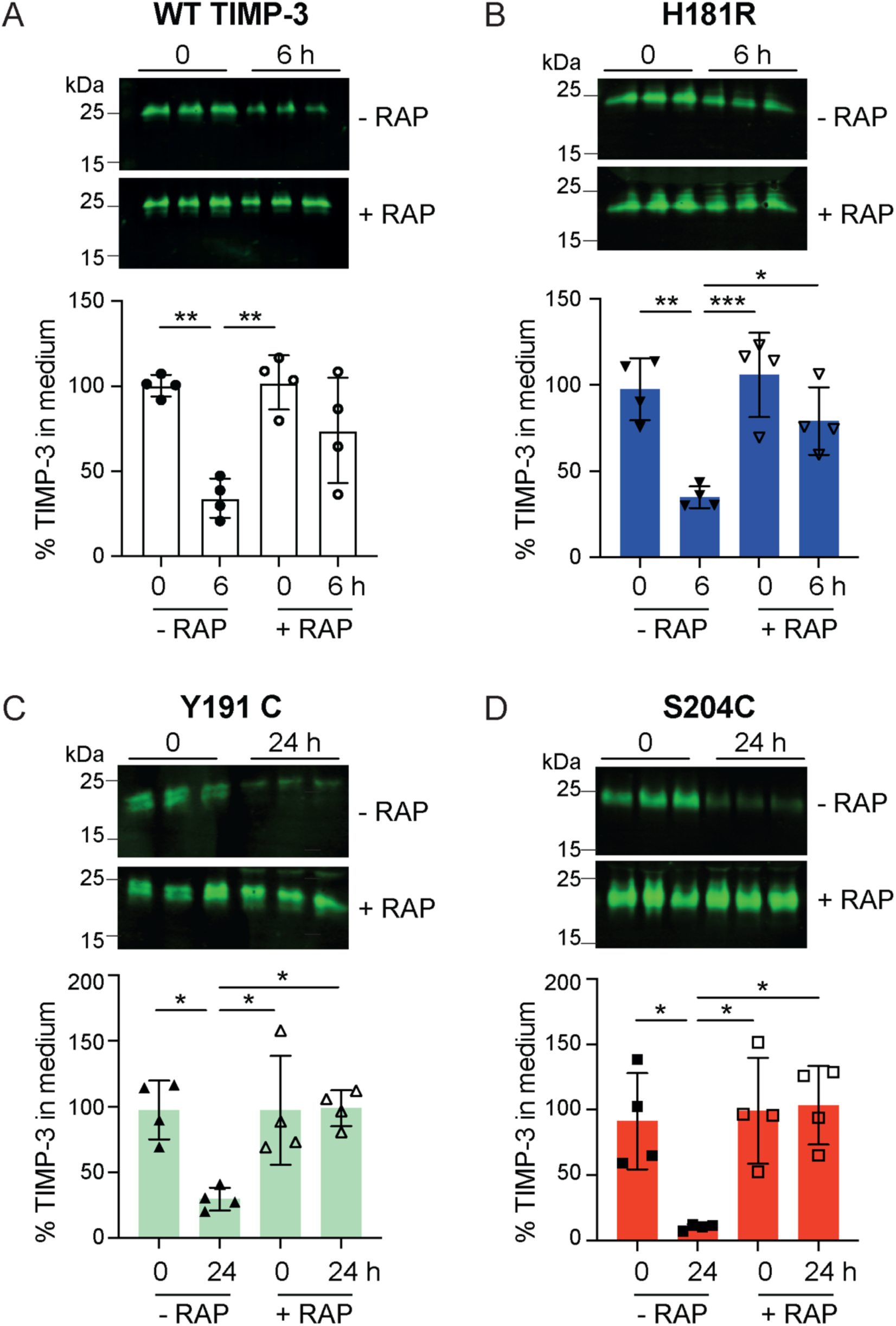
The LRP antagonist RAP blocked clearance of WT and SFD TIMP-3 by ARPE-19 cells. ARPE-19 cells (2.5×10^5^) were pre-treated for 1 h with RAP (0 or 1 μM) before addition of FLAG-tagged TIMP-3 WT (**A**), H181R (**B**), Y191C (**C**) or S204C (**D**) (all 2.5 nM) in DMEM with 0.2% (v/v) FBS (for 0, 6 or 24 h as indicated). Conditioned media were concentrated by TCA precipitation, and TIMP-3 abundance quantified by immunoblotting with anti-FLAG M2 antibody and densitometry, and expressed relative to mean pixel volume at t = 0 h (defined as 100%, n = 4, mean ± SD). Data were assessed for normality by Shapiro-Wilk test, analysed for significance relative to 0 h within each RAP treatment group by two-way ANOVA, and corrected for multiple comparisons with Tukey’s test (* p ≤ 0.05, ** p ≤ 0.01).

### LRP1-dependent and -independent mechanisms of TIMP-3 endocytosis occur

To investigate whether LRP1 was the LRP responsible for TIMP-3 endocytosis, we evaluated several transfection reagents, *LRP1* siRNA sequences and concentrations. In ARPE-19 cells, however, we obtained only 53% knockdown of *LRP1* 24 h after transfection, which fell to 33% knockdown 30 h after transfection (**Fig. 6A-B)**. Despite this low knockdown, an endocytosis assay conducted over the last 6 h of the knockdown period showed that WT TIMP-3 was significantly cleared from conditioned media of cells treated with non-targeting siRNA, but not in those treated with *LRP1* siRNA (**Fig. 6C**), supporting a role for LRP1 in TIMP-3 uptake in these cells. This was supported by ELISA analysis of WT and SFD binding to LRP1, which showed that S204C and Y191C TIMP-3 had lower affinity for LRP1 than WT and H181R TIMP-3 **(Fig. 7A)**.

**Fig. 6:**
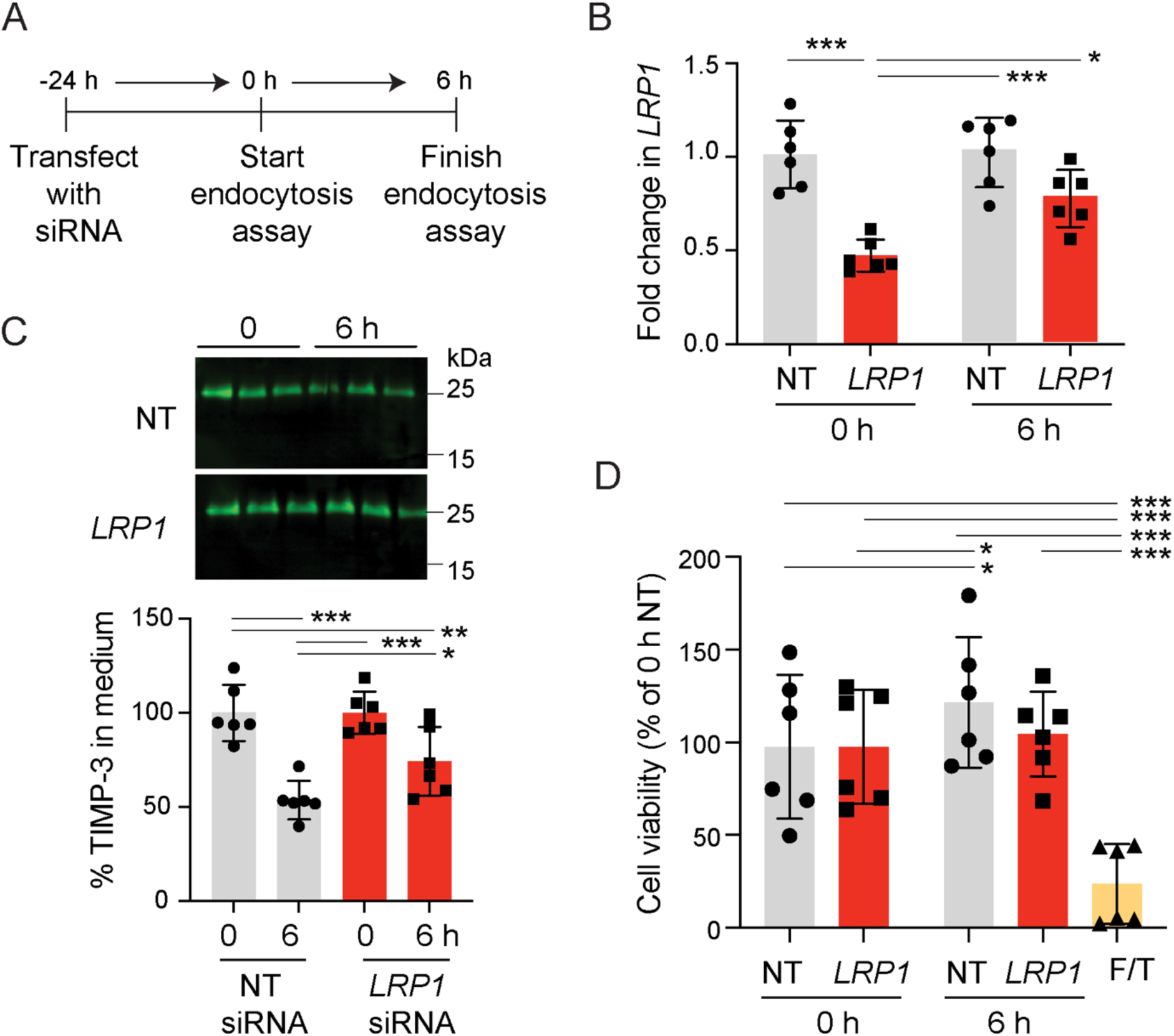
Knockdown of *LRP1* in ARPE-19 cells inhibited clearance of WT TIMP-3. **A**) Schematic representation of experiment, in which ARPE-19 cells (2.5×10^5^) were transfected with non-targeting or LRP1-targeting siRNA (25 nM) 24 h before the start of a 6 h endocytosis assay. Cells were then incubated for 0 or 6 h with FLAG-tagged W TIMP-3 WT (2.5 nM) in DMEM with 0.2% (v/v) FBS. **B**) RT-qPCR used to quantify mRNA expression of *LRP1* relative to *GAPDH* and the NT control at 0 and 6 h (mean ± SD, n = 6). **C**) Conditioned media were concentrated by TCA precipitation, and WT TIMP-3 abundance quantified by immunoblotting with anti-FLAG M2 antibody and densitometry, and expressed relative to mean pixel volume at t = 0 h (defined as 100%, n = 6, mean ± SD). **D**) To assess cell viability, transfected ARPE-19 cells were incubated with MTS reagent (90 min, 37 °C, mean ± SD, n = 6). As a positive control for cell death, cells were freeze-thawed (F/T). Data in **B-D** were assessed for normality by Shapiro-Wilk test, analysed for significance by two-way ANOVA, and corrected for multiple comparisons with Tukey’s test (* p ≤ 0.05, ** p ≤ 0.01, *** p ≤ 0.001).

**Fig. 7:**
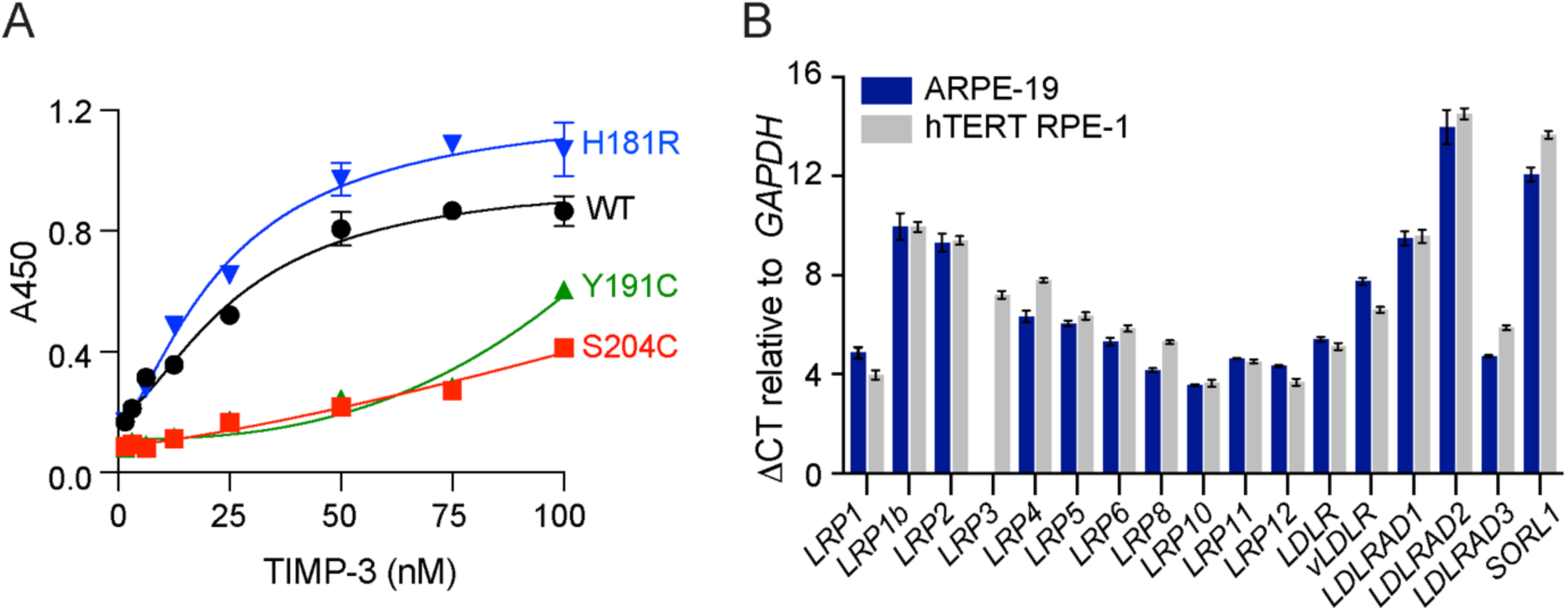
SFD TIMP-3 mutants have reduced affinity for LRP1. **A**) ELISA plates were coated with purified LRP1, and binding of WT, H181R, S204C and Y191C TIMP-3 quantified by ELISA (n = 3, mean ± SD). EC_50_ values for WT (23 ± 1.19 nM) and H181R (22.46 ± 1.20 nM) TIMP-3 were not statistically different from each other, while EC_50_ values for S204C and Y191C were too high to be calculated from these data. **B**) RNA was extracted from ARPE-19 and hTERT RPE-1 cells (5×10^5^) and expression of the indicated *LRP-*related receptors measured by RT-qPCR. Data were normalised to *GAPDH* using the ΔCT method (n = 3, mean ± SD).

In contrast, *LRP1* knockdown in hTERT RPE-1 cells was more effective, reproducible and sustained, with 71% knockdown obtained 24 and 28 h after transfection (**Supplementary Fig. S4A-B**). Significant clearance of WT TIMP-3 was however observed in cells treated with non-targeting as well as *LRP1*-targeting siRNA **Supplementary Fig. S4C**). Taken together with the RAP sensitivity of this cell line (**Supplementary Fig. S3),** this suggests that an LRP other than LRP1 mediates TIMP-3 endocytosis in hTERT RPE-1 cells. RT-qPCR showed that ARPE-19 and hTERT RPE-1 expressed many other LRP family members (Fig. 7B).

Endocytosis of WT and SFD TIMP-3 by an LRP1-null mouse embryonic fibroblast (MEF) cell line (PEA-13)(27) was also investigated. As previously observed (23, 26), WT TIMP-3 was significantly cleared from the conditioned medium of WT but not LRP1-null MEFs (**Fig. 8A**). In contrast, there was no difference in clearance of H181R and Y191C TIMP-3 by the two cell lines (**Fig. 8B-C**), indicating they are endocytosed via an LRP1-independent mechanism in these cells. S204C was also significantly cleared by both cell lines, although uptake by LRP1-null MEFs was significantly lower (**Fig. 8D**).

**Fig. 8:**
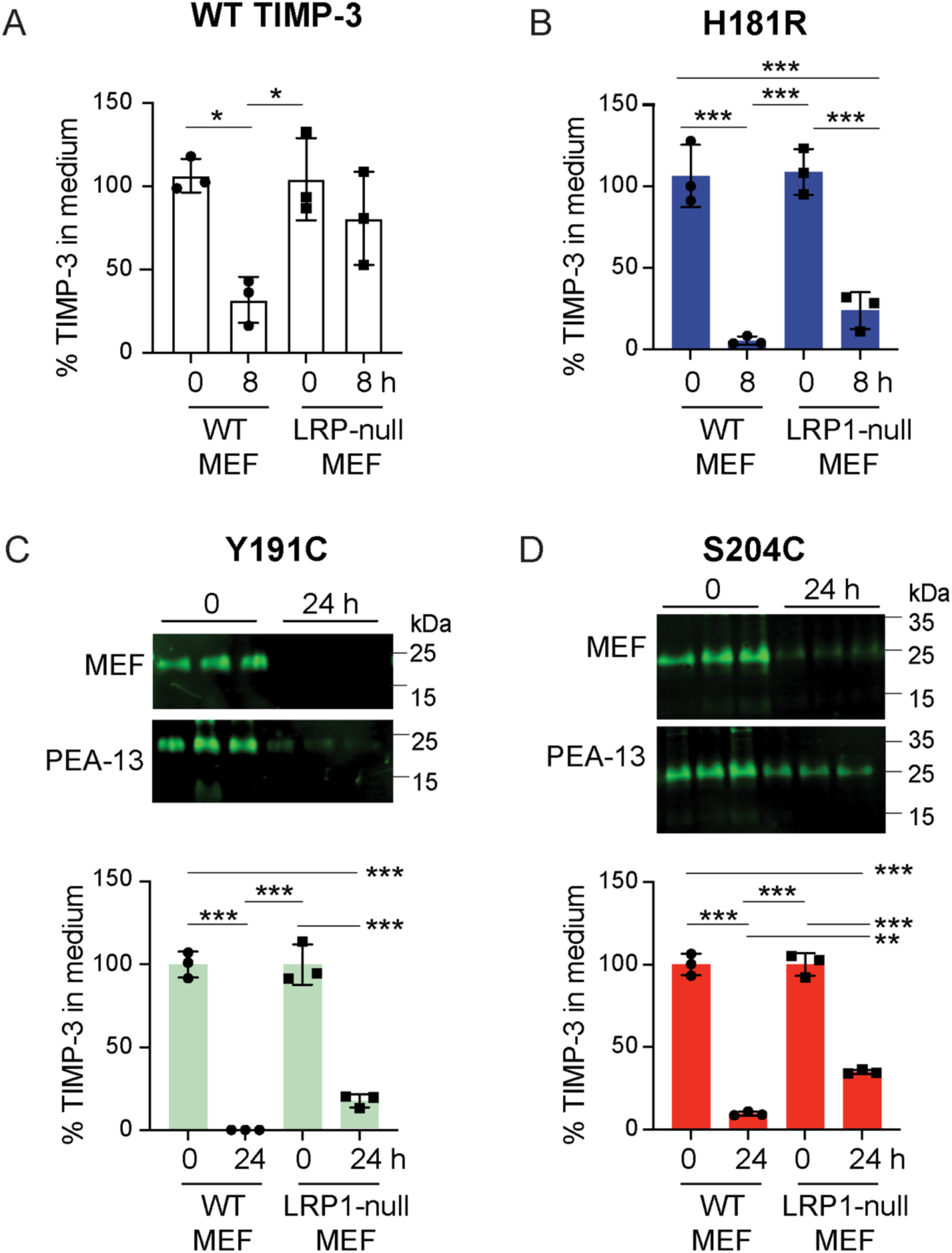
Deletion of LRP1 in mouse embryonic fibroblasts blocked clearance of WT TIMP-3, but not H181R, Y191C or S204C TIMP-3. WT mouse embryonic fibroblasts (WT MEF) and PEA-13 LRP1-null MEFs (5×10^4^) were incubated (0, 8 or 24 h) with FLAG-tagged TIMP-3 WT **(A)**, H181R **(B)**, Y191C **(C)**, or S204C **(D)** (2.5 nM) in DMEM. Conditioned media were harvested at the indicated timepoints, concentrated by TCA precipitation, and TIMP-3 abundance quantified by immunoblotting with anti-FLAG M2 antibody and densitometry. Loss of TIMP-3 variants from conditioned media was calculated relative to total protein staining per lane and to mean pixel volume at t = 0 h (defined as 100%) in each cell type (n = 3, mean ± SD). Data were assessed for normality by Shapiro-Wilk test, analysed for significance with two-way ANOVA, and corrected for multiple comparisons with Tukey’s test (** p ≤ 0.01, *** p ≤ 0.001).

## Discussion

Our previous studies have demonstrated that multiple cell types (fibroblasts, chondrosarcoma cells, chondrocytes, macrophages) endocytose and degrade TIMP-3 via the LRP1 scavenger receptor (22–24). The binding site for LRP1 overlaps with that for HS, so TIMP-3 endocytosis is inhibited by cell surface and extracellular matrix HSPGs (25, 26). Consequently, extracellular TIMP-3 levels are regulated post-translationally by the balance between its interactions with cell surface LRP1 and pericellular HS.

In this study, we show that RPE-derived cell lines widely used in AMD and SFD research (9, 28) also endocytose WT TIMP-3 in an LRP-dependent manner. Endocytosis was inhibited by pre-treatment of ARPE-19 and hTERT RPE-1 cells with RAP, an antagonist of ligand binding to the LRP family of receptors (**Fig. 5**, **Supplementary Fig. S3**). The calculated half-lives of TIMP-3 in these cells (2.6 ± 1.1 h in ARPE-19, and 4.9 ± 2.5 h in hTERT RPE-1) are consistent with clearance rates we observed in other cell types (23, 24, 26). Langton et al. previously calculated a longer half-life (9.5 h) for WT TIMP-3 uptake clearance by ARPE-19 cells (21), but their experiments monitored disappearance of the protein from the extracellular matrix, so binding to HSPGs is likely to have slowed the rate of endocytosis. We found that *LRP1* siRNA knockdown significantly reduced endocytosis in ARPE-19 cells (**Fig. 6**), showing that LRP1 is a key mediator of TIMP-3 endocytosis in these cells. Unexpectedly, *LRP1* knockdown had no effect on endocytosis of WT TIMP-3 by hTERT RPE-1 cells (**Supplementary Fig. S4**), showing for the first time that an LRP other than LRP1 can mediate TIMP-3 endocytosis. It is unclear why hTERT RPE-1 cells differ in this regard, since mRNA expression of *LRP1* is comparable to that in ARPE19 cells (**Fig. 7**). Differences in LRP1 shedding, which can reduce TIMP-3 endocytosis (23), may explain this discrepancy. Alternatively, it may be due to differences in expression of other LRP family members, such as *LRP3*, which is expressed by hTERT RPE-1 but not ARPE-19 cells (**Fig. 7**). Further research is needed to determine which LRP mediates TIMP-3 endocytosis by RPE *in vivo*, and whether this changes with age, disease or inflammation. Notably, *LRP2* was recently identified as a susceptibility locus for AMD in a large, global genome-wide association study (29).

Next, we investigated endocytosis of prototypic SFD variants of TIMP-3, namely S204C and Y191C TIMP-3, as well as H181R TIMP-3, which is unusual among SFD variants in that the mutation does not give rise to an unpaired cysteine residue (30, 9, 4). While all three mutants were endocytosed by ARPE-19 and hTERT RPE-1 cells, S204C and Y191C were endocytosed significantly more slowly than WT TIMP-3 (**Fig. 3** and **Supplementary Fig. S2**). As suggested by Langton et al. (21), the for S204C and S179C TIMP-3, this “[resistance] to turnover” is likely to be the key molecular mechanism leading to the extracellular accumulation of these variants in SFD. The reduced affinity of S204C and Y191C TIMP-3 for LRP1 (**Fig. 7A**), coupled with their RAP sensitivity (**Fig. 5** and **Supplementary Fig. S3**), suggests these mutations hinder LRP-mediated clearance.

Mouse embryonic fibroblasts also endocytosed S204C and Y191C TIMP-3 more slowly than WT TIMP-3, but deletion of LRP1 had no effect (for Y191C) or a limited effect (for S204C) on this slower endocytosis (**Fig. 8**), indicating that an LRP1-independent route can mediate slow clearance of these mutants in some cell types. This may be mediated by another LRP or by another mechanism. ARPE-19 and hTERT RPE-1 cells appear to lack this LRP-independent clearance route, indicating that the retina may be particularly sensitive to SFD TIMP-3 mutations. Meunier et al. (31) report lung pathology in SFD patients with the Y191C and S38C variants of TIMP-3, raising the possibility that this may be another site where LRP-independent clearance mechanisms are limited, at least for some SFD variants.

In contrast to S204C and Y191C TIMP-3, H181R TIMP-3 exhibited endocytosis kinetics indistinguishable from WT TIMP-3, suggesting it does not accumulate extracellularly due to impaired endocytosis. Instead, the H181R mutation may increase affinity for HS, given that the mutation introduces a more basic residue near the positively-charged face of TIMP-3 that interacts with HS (32). Alternatively, this could change the specificity of HS binding as has been seen previously for the Y402H polymorphism in complement Factor H (33–35), that is associated with increased risk of AMD (36–38). This could subtly alter post-translational trafficking, leading to a slow extracellular accumulation of the mutant over the 50 or so years before onset of clinical symptoms in SFD patients carrying this variant. The E162K variant of TIMP-3 also introduces an additional basic residue to TIMP-3, and may behave similarly.

The heterogeneity in molecular mechanisms of SFD TIMP-3 accumulation suggests that therapeutic strategies to enhance TIMP-3 clearance may be more effective for some SFD variants than others. For example, clearance of H181R TIMP-3 could be increased by solubilising it from the extracellular matrix with sulfated glycans such as heparin, but such an approach would be less effective for variants with reduced LRP affinity. This underscores the importance of genotyping SFD patients to tailor personalised therapeutic interventions.

The reduced turnover of SFD TIMP-3 has been attributed to their tendency to form dimers and multimers (21). Most SFD mutations give rise to unpaired cysteine residues that cause multimerization via aberrant inter-molecular disulfide bond formation, but even variants that do not give rise to unpaired cysteine residues (e. g. H181R and E162K) have been shown to form dimers (17, 39) through an unknown molecular mechanism. In our experiments, low levels of multimeric SFD TIMP-3 were detected in conditioned media of transfected HEK-293-EBNA cells, but these higher molecular weight species were largely lost during purification by M2 anti-FLAG affinity chromatograph, presumably due to aggregation and non-specific retention on the resin. Thus, our analysis reflects rates of monomeric TIMP-3 uptake, demonstrating that even in their monomeric forms, S204C and Y191C TIMP-3 exhibit reduced LRP binding and endocytosis. LRP ligands, including TIMP-3, bind to the receptors via pairs of basic residues separated by 21 angstroms (26, 40, 41), and SFD mutations may alter TIMP-3 conformation in a way that impairs this interaction. Further studies are required to investigate rates of endocytosis for multimeric SFD species. This could be done using crude media from transfected HEK-293-EBNA cells, but these cells also express LRP1, which may be shed into media and act as a decoy receptor (23). Transient transfection of RPE cells with WT and SFD could also be used to compare rates of endocytosis, although in our hands, expression efficiency of SFD variants in ARPE-19 cells was several folds lower than that of WT TIMP-3, making it difficult to accurately compare their rates of uptake. Competition for TIMP-3 binding by pericellular HSPGs could be controlled for by treatment of cells with sodium chlorate (42).

Beyond SFD, our findings have implications for AMD. TIMP-3 is a component of AMD drusen deposits (7, 43), and genome-wide association studies have identified AMD risk loci near the *TIMP3* gene (38, 44). As with SFD variants of TIMP-3 (20), levels of WT TIMP-3 in the Bruch’s membrane increases with age (7, 45) without a corresponding increase in mRNA expression (46), indicating that its abundance is primarily regulated post-translationally. TIMP-3 levels in the Bruch’s membrane also increase in AMD in people with the high-risk chromosome 1 haplotype (6). Several other drusen components (e.g. clusterin, vitronectin) and proteins encoded by AMD risk loci (e.g. HTRA1, APOE, ADAMTS9, C3, LIPC, MMP-9, VEGFA) also interact with LRP1 (47–55) as well as heparin or HS (56, 57), suggesting that disruption of the equilibrium between LRP1 and HS may drive drusen formation. Age-related and inflammation-dependent changes in LRP1 expression (58–61) and HS structure (62–64) may further influence this equilibrium.

Understanding how SFD mutations disrupt extracellular TIMP-3 trafficking will not only clarify disease mechanisms but may also provide insights into broader physiological and pathological processes involving TIMP-3, such as development, wound healing, osteoarthritis and chronic inflammation. Future work should explore therapeutic strategies to restore proper TIMP-3 clearance, potentially benefiting a range of age-related and degenerative conditions.

## Materials and Methods

### Materials

Brij 35 (30% w/v solution), calcium chloride (CaCl_2_), dimethyl sulfoxide, glycerol, Gibco cell dissociation solution, Gibco DMEM (4.5 g/L D-Glucose, L-glutamine, no pyruvate), Gibco Opti-MEM, hygromycin B solution, phosphate-buffered saline, sodium azide (NaN_3_), sodium chloride, sodium chlorate, sodium dodecyl sulfate (SDS), sodium pyruvate, Tris base, trypsin-EDTA (0.25% w/v), and Tween 20 were from ThermoFisher Scientific (Loughborough, UK). Heat-inactivated foetal bovine serum, penicillin/streptomycin were from Merck (Gillingham, UK). Lipofectamine LTX and Lipofectamine 3000 transfection reagents were from ThermoFisher. Human MMP-1 (65) and receptor-associated protein (RAP) (61) were expressed and purified as described previously.

### Cell culture

HEK-293-EBNA, ARPE-19, hTERT RPE-1, PEA-13 and MEF cells were from ATCC, Manassas, VA. All cells were maintained in DMEM (defined as including 100 units/ml penicillin, 100 units/ml streptomycin, and 1 mM sodium pyruvate) supplemented with 10% (v/v) FBS at 37 °C in 5% (v/v) CO_2_. For some experiments, cells were cultured in DMEM without FBS, referred to as serum-free DMEM.

### Generation of TIMP-3 mutants

TIMP-3 mutants were generated as described previously, using the QuikChange II XL site-directed mutagenesis kit (Agilent Technologies, Cheshire, UK), with C-terminally FLAG-tagged WT TIMP-3 in pCEP4 mammalian expression vector as the template (26). Primers (Merck, Gillingham, UK) for generation of H181R were 5’-ACC CTG GCT ACC AGT CCA AAC GCT ACG CCT GCA TCC GGC AGA AGG G-3’ (forward) and 5’-CCG CCC TTC TGC CGG ATG CAG GCG TAG CGT TTG GAC TGG TAG CAG GG-3’ (reverse). Primers for generation of S204C were 5’-AGG ATG GGC CCC CCC GGA TAA ATG CAT CAT CAA TGC CAC AGA CCC CG-3’ (forward) and 5’-CCT ACC CGG GGG GGC CTA TTT ACG TAG TAG TTA CGG TGT CTG GG-3’ (reverse). Primers for generation of Y191C were 5’-TGC ATC CGG CAG AAG GGC GGC TGC TGC AGC TGG TAC CGA GGA TGG G-3’ (forward) and 5’-CGG ACG TAG GCC GTC TTC CCG CCG ACG ACG TCG ACC ATG GCT CCT AC-3’ (reverse). Calculated melting temperatures ranged between 91.2 °C and 95.5 °C, and guanine/cytosine-content between 44 and 48%. PCR conditions were as per the manufacturer’s instructions, except that the extension time per cycle was increased to 150 s/kb (1435 s). PCR products were digested with *Dpn*I and transformed into *E. coli* XL10-Gold ultracompetent bacteria by heat shock at 42 °C before plating on agar plates containing 100 μg/ml carbenicillin. Candidate colonies were screened by DNA sequencing (Genewiz from Azenta Life Sciences, Bishop’s Stortford, UK) using the pCEP4 forward primer 5’-GGA GGT CTA TAT AAG CAG AGC TCG-3’ and reverse primer 5’-CTG CAT TCT AGT TGT GGT TTG TCC-3’.

### Purification of WT and mutant TIMP-3

C-terminally FLAG-tagged WT and mutant human TIMP-3 were expressed in HEK-293-EBNA cells and purified by anti-FLAG M2 affinity chromatography (66). HEK-293-EBNA cells were transfected with pCEP4 expression plasmids encoding WT or mutant TIMP-3 using Lipofectamine 3000 and selected using hygromycin B (200 μg/ml). After expansion of cell number, transfected cells were cultured in serum-free DMEM containing 30 mM sodium chlorate (NaClO_3_) for 48 h. Conditioned media were harvested, centrifuged to remove cell debris, and sequentially filtered through 40 μm (Greiner Bio-One, Kremsmünster, Austria) and 0.22 μm filters (VWR, Lutterworth, UK). Filtered media were passed over an anti-FLAG M2 agarose resin (Merck, Gillingham, UK) equilibrated in either serum-free DMEM or TNC buffer [50 mM Tris.HCl, pH 7.5, 150 mM NaCl, 10 mM CaCl_2_, 0.02% (w/v) NaN_3_, 0.05% (w/v) Brij 35]. The resin was washed in equilibration solution containing 600 mM NaCl to remove non-specifically bound proteins, and returned to equilibration solution. Bound WT or mutant TIMP-3 were eluted from the resin with FLAG peptide (200 μg/ml, Cambridge Biosciences, Cambridge, UK) in equilibration solution and stored at −80 °C in Eppendorf LoBind microcentrifuge tubes (ThermoFisher Scientific, Loughborough, UK). The purity of isolated TIMPs was confirmed by silver staining (Pierce Silver Stain Kit, ThermoFisher Scientific, Loughborough, UK) and their active concentrations determined by titration against a known concentration of human MMP-1 as described previously (26, 66).

### SDS-PAGE and immunoblotting

Protein samples were mixed with non-reducing SDS-PAGE sample buffer [100 mM Tris.HCl, pH 6.8, 2% (w/v) SDS, 15% (v/v) glycerol, 0.01% (w/v) bromophenol blue)] and electrophoresed on 4-15% precast Mini-PROTEAN TGX Gels (Bio-Rad, Watford, UK), or 12 and 15% Tris-glycine polyacrylamide gels polymerised in-house from 30% (w/v) acrylamide (37.5:1 acrylamide:bisacrylamide ratio, Severn Biotech, Kidderminster, UK). PageRuler Plus prestained protein ladder and all other electrophoresis reagents were from ThermoFisher Scientific (Loughborough, UK).

In some cases, conditioned media were concentrated before electrophoresis by addition of trichloroacetic acid (TCA, 6.1 N, ThermoFisher Scientific, Loughborough, UK) to 5% (v/v). After overnight incubation at 4 °C, solutions were centrifuged (16 000 RCF, 10 min, 4 °C) and pellets resuspended in SDS-PAGE sample buffer.

For analysis of disulfide bond formation, samples were reduced with β-mercaptoethanol (20 mM, 45 min, 56 °C), cooled (15 min, 20 °C), and alkylated with iodoacetamide (20 mM, 30 min, 20 °C) before addition of SDS-PAGE sample buffer. For silver staining, a Pierce silver stain kit from ThermoFisher Scientific (Loughborough, UK) was used.

For immunoblotting, gels were blotted to low fluorescence Immobilon-FL PVDF membranes (Merck, Gillingham, UK) using the Trans-Blot Turbo Transfer System (Bio-Rad, Watford, UK). Blots were stained for total protein using Revert Total Protein Stain (LI-COR, Cambridge, UK), imaged using an Odyssey CLx imager (LI-COR, Cambridge, UK), and destained following the manufacturer’s protocol. Blots were incubated in Intercept blocking buffer (1 h, 20 °C, LiCOR, Cambridge, UK), and then with the required primary antibody in 50% (v/v) Intercept Blocking Buffer supplemented with 0.2% (v/v) Tween 20 (overnight, 4 °C). Membranes were washed [4x5 min in TBS with 0.1% (v/v) Tween 20], and incubated with an appropriate secondary antibody diluted in 50% (v/v) Intercept Blocking Buffer supplemented with 0.2% (v/v) Tween 20 and 0.2% (w/v) SDS (1-2 h, 20 °C). After washing [4x5 min in TBS with 0.1% (v/v) Tween 20], blots were imaged and quantified using an Odyssey CLx imager. Bands of interest were quantified using Image Studio Version 4.0.21 and normalised to total protein signal in each lane.

Anti-FLAG M2 antibody was from Merck (Gillingham, UK). IRDye 800CW goat anti-mouse secondary antibody was from LI-COR (Cambridge, UK).

### MTS assay for cell viability

Cell viability was assessed using the CellTiter 96 Aqueous One Solution Cell Proliferation Assay (Promega, Chilworth, UK) following the manufacturer’s instructions. Briefly, cells were collected in cell dissociation solution and resuspended in serum-free DMEM. MTS reagent was added to cells for 90 min [37 °C, 5% (v/v) CO_2_] and absorbance at 490 nm measured in a SPECTROstar Omega spectrophotometer (BMG LABTECH, Aylesbury, UK). As a control, cells were repeatedly frozen at −80 °C and thawed before quantification. Absorbance was converted to cell number using a standard curve prepared by plating a range of known cell numbers.

### Flow cytometry analysis of apoptosis

ARPE-19 cells (2.5×10^5^) were plated overnight in DMEM with 10% (v/v) FBS, washed into serum-free DMEM, and incubated with WT, H181R or S204C TIMP-3 (100 nM in serum-free DMEM) for 16 h [37 °C, 5% (v/v) CO_2_]. A positive control for cell death was generated by repeated freeze-thawing between −80 °C and room temperature. Cells were trysinised and resuspended in Annexin V Binding Buffer (50 µl, BioLegend, London, UK). After addition of FITC-Annexin V (1 µl, BioLegend, London, UK) and propidium iodide (0.33 µl, BioLegend, London, UK), cells were vortexed gently and incubated for 15 min at 4 °C before analysis (5 000 events for freeze/thaw and 20 000 for other conditions) on a FACSymphony A1 Cell Analyzer (BD Biosciences, Wokingham, UK). Calibration was performed using Cytometer Setup and Tracking Research Beads (BD Biosciences, Wokingham, UK). Compensations and Fluorescence Minus One controls were performed to exclude auto- and background fluorescence. Data were analysed using FloJo v10.10.0 Software (BD Life Sciences, Ashland, USA).

### Endocytosis assays

ARPE-19 (2.5×10^5^), hTERT RPE-1 cells (2.5×10^5^), WT MEFs (5×10^5^) or LRP1-null PEA-13 MEFs (5×10^5^) (27) were plated overnight in DMEM with 10% (v/v) FBS, washed into DMEM containing 0.2% (v/v) FBS, and incubated with WT, H181R or S204C TIMP-3 (2.5 nM in TNC buffer) for 0 - 24 h [37 °C, 5% (v/v) CO_2_]. Conditioned media were harvested at the indicated timepoints, concentrated by TCA precipitation, and electrophoresed under non-reducing conditions. Gels were blotted to low fluorescence Immobilon-FL PVDF membranes, stained for total protein, and analysed using anti-FLAG M2 primary antibody with IRDye 800CW goat anti-mouse secondary antibody. Loss of TIMP-3 variants from conditioned media was calculated relative to total protein staining per lane and to pixel volume at t = 0 h (defined as 100%, n = 3, mean ± SD). Data were fitted to a one-phase exponential decay model to calculate half-lives and the percentage of each TIMP-3 variant remaining in conditioned media at 6 h.

### Immunofluorescent microscopy

Glass coverslips were coated in 10 mg/ml gelatin dissolved in H_2_O (15 min, 20 °C) before fixation in 1% (w/v) glutaraldehyde in TBS (20 min, 20 °C) and quenching in 1 M ammonium chloride (15 min, 20 °C). After washing the coverslips in DMEM, ARPE-19 cells (1×10^5^) were seeded and left to adhere overnight. Cells were washed 3 times in serum-free DMEM, and incubated [37 °C, 5% (v/v) CO_2_] with WT or SFD TIMP-3 (40 nM) in DMEM with 0.2% (v/v) FBS for 2 h (for WT and H181R TIMP-3) or 18 h (for Y191C and S204C TIMP-3). After this incubation, the cells were washed in PBS and fixed in 3% (v/v) paraformaldehyde (Merck, Gillingham, UK) in PBS (10 min, 25 °C), followed by a further wash in PBS and blocking in PBS containing 5% (v/v) goat serum (ThermoFisher Scientific, Loughborough, UK) and 3% (v/v) FBS (1 h, 20 °C). Cells were permeabilised with PBS containing 0.1% (v/v) Triton X-100 (15 min, 20 °C) and incubated with anti-FLAG M2 primary antibody (Merck, Gillingham, UK) in blocking solution (3 h, 25 °C). After washing 3 times in PBS, cells were incubated with an Alexa Fluor-488-conjugated goat anti-mouse IgG secondary antibody (Invitrogen, via ThermoFisher Scientific, Loughborough, UK) to visualise FLAG staining. Actin was stained with Alexa Fluor-647-conjugated phalloidin (Invitrogen, via ThermoFisher Scientific, Loughborough, UK), and nuclei visualised with DAPI staining (Invitrogen, via ThermoFisher Scientific, Loughborough, UK). Cells were visualised using a Zeiss LSM 800 laser scanning confocal microscope with a 60x objective lens.

### siRNA knockdown

ARPE-19 or hTERT RPE-1 cells (2.5×10^5^) were plated overnight in DMEM with 10% (v/v) FBS, and washed three times in serum-free DMEM before transfection in Opti-MEM containing Lipofectamine LTX and either non-targeting (Dharmacon ON-TARGETplus non-targeting pool, Horizon Discovery, Cambridge, UK) or *LRP1*-targeting (Dharmacon ON-TARGETplus SMARTpool catalogue number L-004721-00-005, Horizon Discovery, Cambridge, UK) siRNA. After 6 h, media were changed to DMEM with 10% (v/v) FBS and cells incubated for a further 18 h before endocytosis assays were started.

### RT-qPCR

RNA was isolated using RNeasy RNA extraction kits (QIAGEN, Manchester, UK) and quantified using a Nanodrop One (ThermoFisher Scientific, Loughborough, UK) before reverse transcription (400-500 ng of RNA) using an Applied Biosystems high-capacity cDNA synthesis kit (ThermoFisher Scientific, Loughborough, UK)). After dilution 1:4 with RNAse-free water (QIAGEN, Manchester, UK), cDNA was quantified by qPCR on a QuantStudio 7 Flex system (Applied Biosystems, Waltham, USA) using SYBR-Green master mix (PCR Biosystems, London, UK) and KiCqStart SYBR Green Primers (1 µM final concentration of forward and reverse primers, Merck, Gillingham, UK). Sequences of primers used is given in **Supplementary Table 1**. Melt curves were analysed on each run to ensure that only a single species of PCR product was generated. Changes in expression of genes of interest were calculated using the ΔΔCT method and expressed relative to *GAPDH*.

### LRP1 ELISA

Binding of WT and SFD TIMP-3 to LRP1 was quantified by ELISA as previously described (26). In brief, medium-binding ELISA plates (Greiner Bio-One, Stonehouse, UK) were coated (overnight, 4 °C) with isolated human LRP1 (5 nM; BioMac, Leipzig, Germany) and blocked with 3% (w/v) BSA in Tris-buffered saline (TBS, 50 mM Tris.HCl, pH 7.5, 115 mM NaCl). Wells were washed with TBS containing 0.1% (v/v) Tween 20 after this and every subsequent step. WT or mutant TIMP-3 (1.5–100 nM) in blocking solution was added (3 h, 37 °C), and binding to LRP1 detected with anti-FLAG M2 primary antibody and anti-mouse-HRP-conjugated secondary antibody in the same buffer. Absorbance at 450 nm was measured using a FLUOstar OMEGA microplate reader (BMG Labtech).

### Statistical analysis

All data were assessed for normality with Shapiro-Wilks test, and analysed for significance with corrections for multiple comparisons as indicated in the figure legends using GraphPad Prism v10.4.1 (GraphPad Software, Boston, USA). All significant differences are indicated on the figures.

### Funding and additional information

JHJB and LT were supported by The Macular Society. KH was supported by the UKRI BBSRC Norwich Research Park Bioscience Doctoral Training Programme BB/T008717/1

## Abbreviations

AMD: age-related macular degeneration
DMEM: Dulbecco’s modified Eagle Medium
ECM: extracellular matrix
FBS: foetal bovine serum
HS: heparan sulfate
HSPG: heparan sulfate proteoglycan
LRP: low-density lipoproteinreceptor-related protein
MMP-1: matrix metalloproteinase 1
RAP: receptor-associated protein
RPE: retinal pigment epithelium
SDS: sodium dodecyl sulfate
SFD: Sorsby fundus dystrophy
TBS: Tris-buffered saline
TNC: Tris-sodium-calcium buffer
TIMP-3: tissue inhibitor of metalloproteinases 3
WT: wild-type.

## Figure Legends

**Supplementary Fig. S1:**
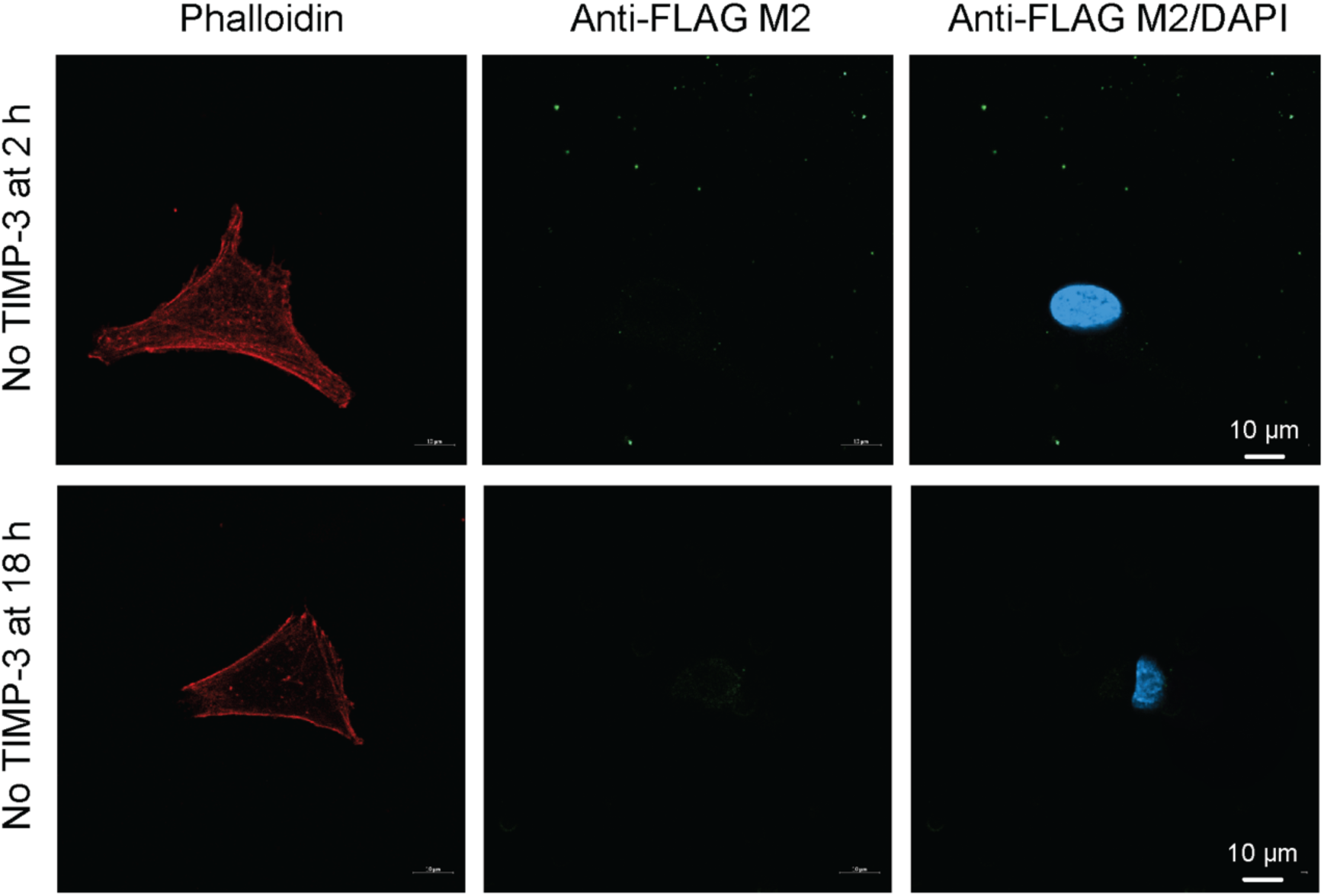
Negative controls for fluorescent microscopy. ARPE-19 cells (1×10^5^) were seeded on gelatin-coated coverslips, and incubated in DMEM with 0.2% (v/v) FBS for either 2 h (upper panels) or 18 h (lower panels). Cells were permeabilised and stained with anti-FLAG M2 primary antibody and Alexa Fluor-488-conjugated goat anti-mouse IgG. The actin cytoskeleton was stained with Alexa Fluor-647-conjugated phalloidin and nuclei stained with DAPI.

**Supplementary Fig. S2:**
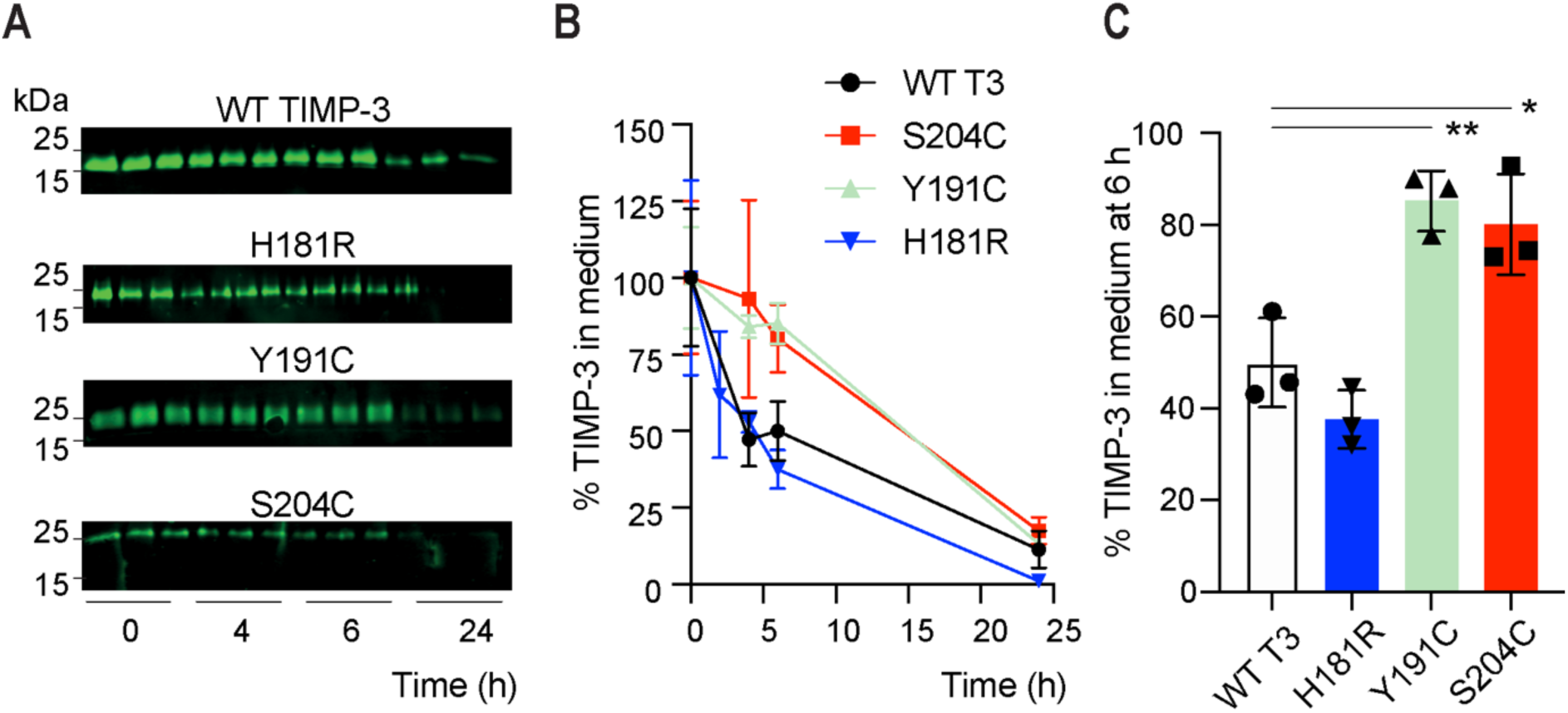
hTERT RPE-1 cells endocytosed WT TIMP-3 and H181R more rapidly than the S204C and Y191C variants. hTERT RPE-1 cells (2.5×10^5^) were incubated (0-24 h) with FLAG-tagged TIMP-3 WT, H181R, Y191C or S204C (2.5 nM) in DMEM with 0.2% (v/v) FBS. **A**) Conditioned media were harvested at the indicated timepoints, concentrated by TCA precipitation, and TIMP-3 abundance quantified by immunoblotting with anti-FLAG M2 antibody and densitometry. Representative immunoblots are shown. **B**) Loss of TIMP-3 variants from conditioned media was calculated relative to total protein staining per lane and to pixel volume at t = 0 h (defined as 100%, n = 3, mean ± SD). **C**) The percentage of each TIMP-3 variant remaining in conditioned media at 6 h was calculated, with data assessed for normality by Shapiro-Wilk test, analysed for significance relative to WT TIMP-3 with one-way ANOVA, and corrected for multiple comparisons with Tukey’s test (* p ≤ 0.05, ** p ≥ 0.01).

**Supplementary Fig. S3:**
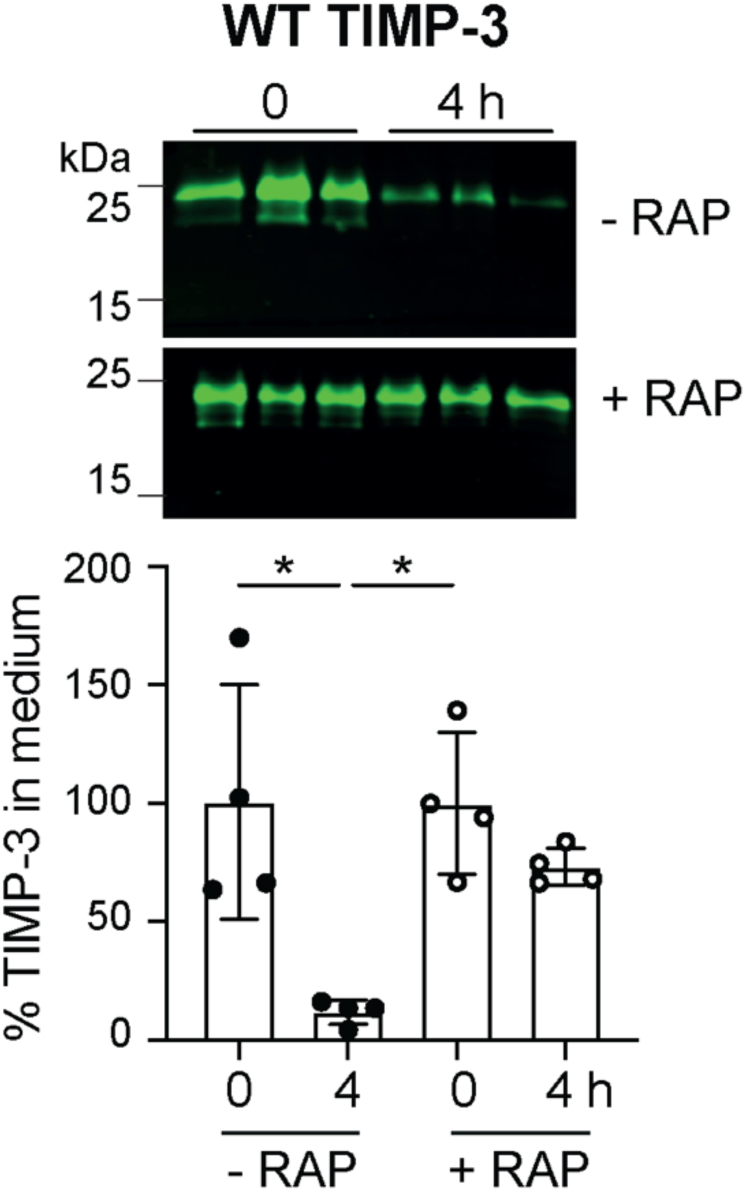
The LRP antagonist RAP blocked clearance of WT TIMP-3 by hTERT RPE-1 cells. hTERT RPE-1 cells (2.5×10^5^) were pre-treated for 1 h with RAP (0 or 1 μM) before addition of FLAG-tagged TIMP-3 WT (2.5 nM) in DMEM with 0.2% (v/v) FBS (for 0 or 4 h as indicated). Conditioned media were concentrated by TCA precipitation, and TIMP-3 abundance quantified by immunoblotting with anti-FLAG M2 antibody and densitometry, and expressed relative to mean pixel volume at t = 0 h (defined as 100%, n = 4, mean ± SD). Data were assessed for normality by Shapiro-Wilk test, analysed for significance by two-way ANOVA, and corrected for multiple comparisons with Tukey’s test (* p ≤ 0.05).

**Supplementary Fig. S4:**
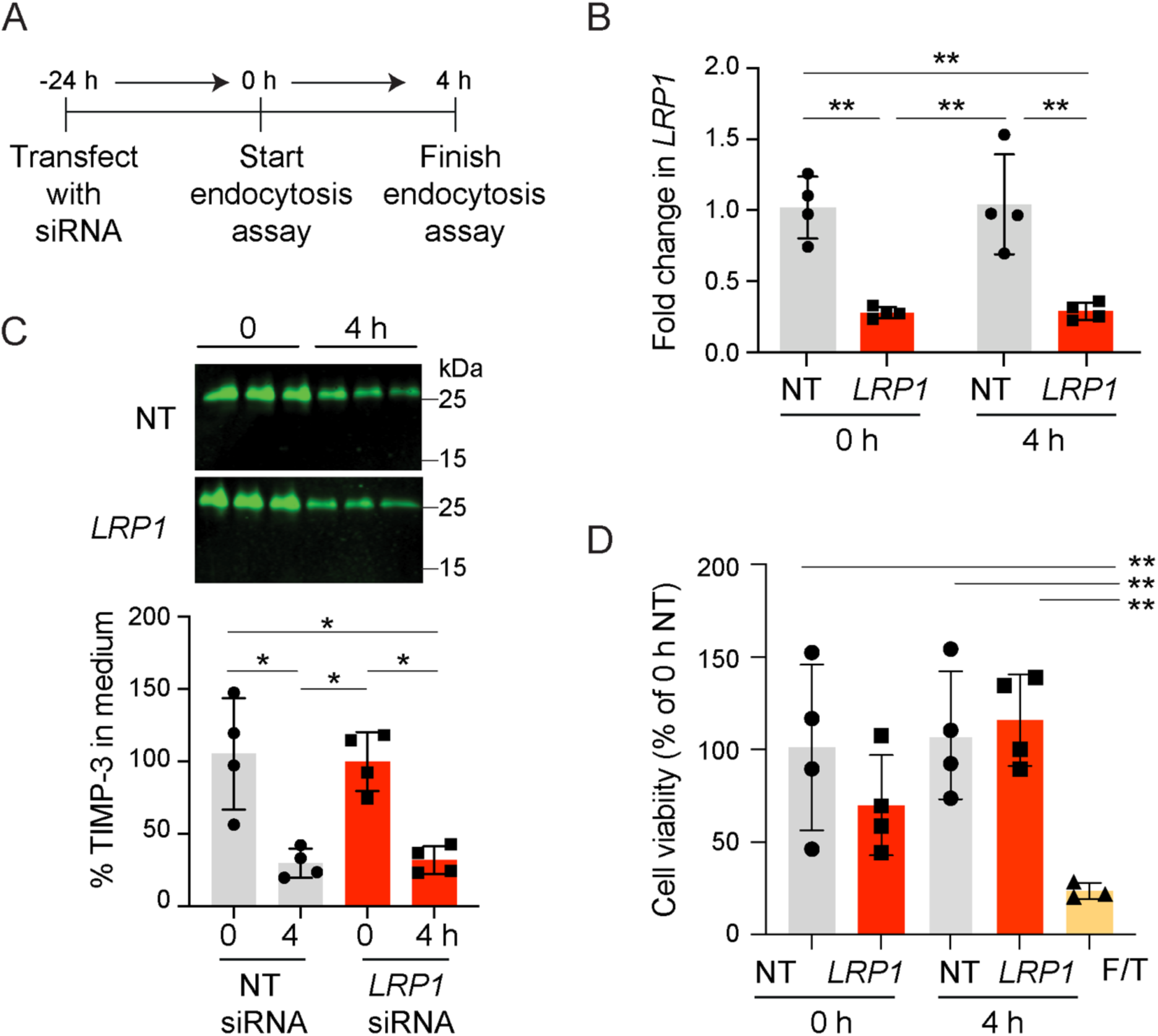
Knockdown of *LRP1* in hTERT RPE-1 cells did not alter clearance of WT TIMP-3. **A**) Schematic representation of experiment, in which hTERT RPE-1cells (2.5×10^5^) were transfected with non-targeting or LRP1-targeting siRNA (25 nM) 24 h before the start of a 4 h endocytosis assay. Cells were then incubated for 0 or 4 h with FLAG-tagged WT TIMP-3 (2.5 nM) in DMEM with 0.2% (v/v) FBS. **B)** RT-qPCR used to quantify mRNA expression of *LRP1* relative to *GAPDH* and the NT control at 0 and 4 h (mean ± SD, n = 4). **C**) Conditioned media were concentrated by TCA precipitation, and WT TIMP-3 abundance quantified by immunoblotting with anti-FLAG M2 antibody and densitometry, and expressed relative to mean pixel volume at t = 0 h (defined as 100%, n = 4, mean ± SD). **D**) To assess cell viability, transfected hTERT RPE-1 cells were incubated with MTS reagent (90 min, 37 °C, mean ± SD, n = 3-4). As a positive control for cell death, cells were freeze-thawed (F/T). Data in **B-D** were assessed for normality by Shapiro-Wilk test, and analysed for significance by two-way ANOVA, and corrected for multiple comparisons with Tukey’s test (* p ≤ 0.05, ** p ≤ 0.01).

**Supplementary Table 1:**
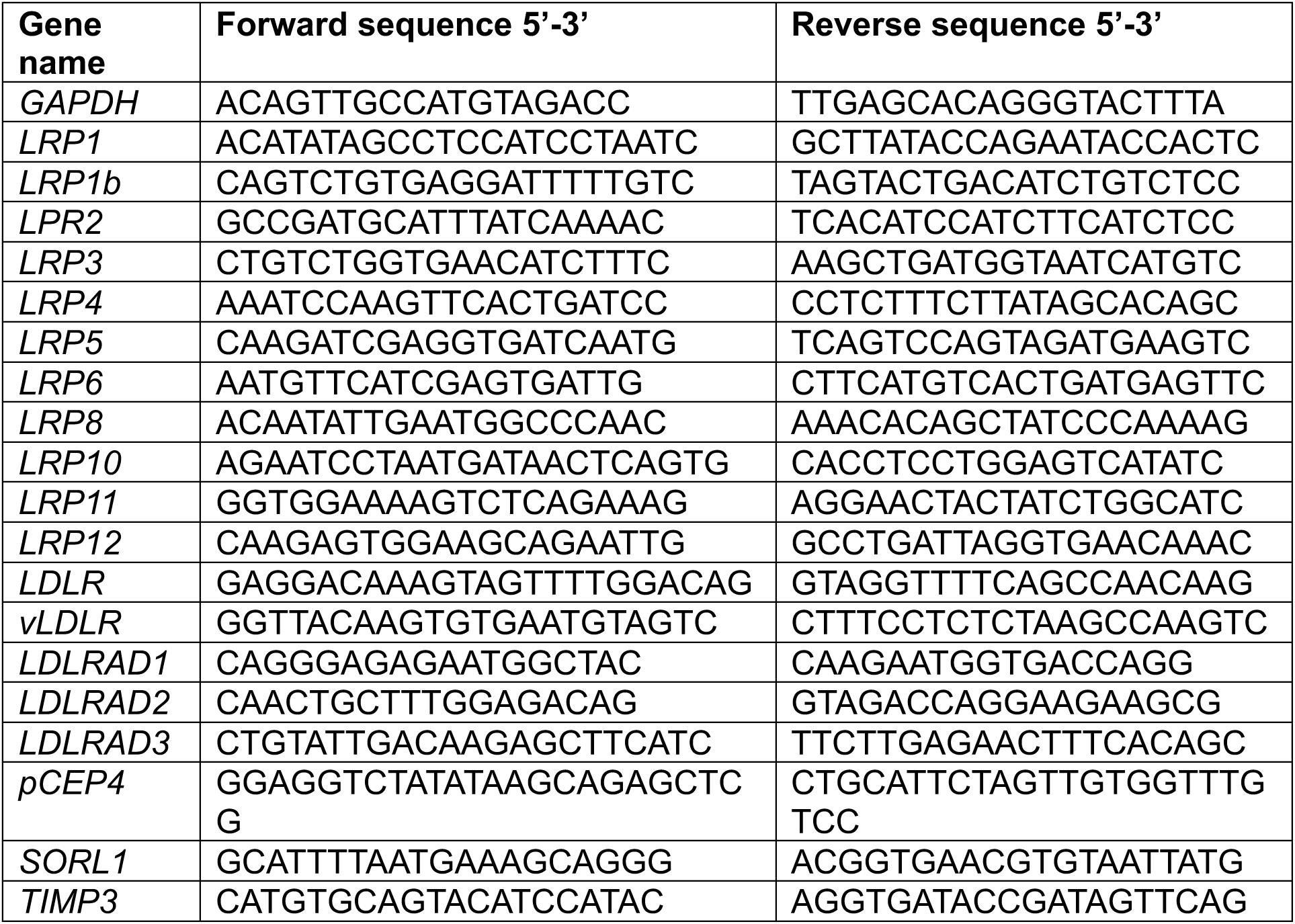
KiCqStart SYBR Green DNA primers (Merck, Gillingham, UK) used for RT-qPCR.

